# Leveraging Multi-Echo EPI to Enhance BOLD Sensitivity in Task-based Olfactory fMRI

**DOI:** 10.1101/2024.01.15.575530

**Authors:** Ludwig Sichen Zhao, Clara U. Raithel, M. Dylan Tisdall, John A. Detre, Jay A. Gottfried

## Abstract

Functional magnetic resonance imaging (fMRI) using blood-oxygenation-level-dependent (BOLD) contrast relies on gradient echo echo-planar imaging (GE-EPI) to quantify dynamic susceptibility changes associated with the hemodynamic response to neural activity. However, acquiring BOLD fMRI in human olfactory regions is particularly challenging due to their proximity to the sinuses where large susceptibility gradients induce magnetic field distortions. BOLD fMRI of the human olfactory system is further complicated by respiratory artifacts that are highly correlated with event onsets in olfactory tasks.

Multi-echo EPI (ME-EPI) acquires gradient echo data at multiple echo times (TEs) during a single acquisition and can leverage signal evolution over the multiple echo times to enhance BOLD sensitivity and reduce artifactual signal contributions. In the current study, we developed a ME-EPI acquisition protocol for olfactory task-based fMRI and demonstrated significant improvement in BOLD signal sensitivity over conventional single-echo EPI (1E-EPI). The observed improvement arose from both an increase in BOLD signal changes through a *T_2_\**-weighted echo combination and a reduction in non-BOLD artifacts through the application of the Multi-Echo Independent Components Analysis (ME-ICA) denoising method. This study represents one of the first direct comparisons between 1E-EPI and ME-EPI in high-susceptibility regions and provides compelling evidence in favor of using ME-EPI for future task-based fMRI studies.

## 1 Introduction

Functional magnetic resonance imaging (fMRI) is one of the most common imaging techniques used to non-invasively study both localized neural activation and whole-brain functional connectivity in humans. Blood-oxygenation-level-dependent (BOLD) fMRI measures brain activity by detecting local magnetic susceptibility fluctuations caused by changes in deoxyhemoglobin concentration. Gradient echo echo-planar imaging (GE-EPI) is the dominant BOLD imaging strategy due to its high temporal signal-to-noise ratio and rapid acquisition capabilities (Budde et al., 2014). However, artifacts caused by the local inhomogeneous magnetic field surrounding high-susceptibility regions present a major technical challenge in utilizing GE-EPI to measure BOLD signals. Olfactory regions, including the orbitofrontal cortex (OFC), amygdala, piriform (olfactory) cortex, and entorhinal cortex are particularly affected by susceptibility-induced artifacts and signal drop-out due their location near the ethmoid sinuses (air-filled cavities in the bone surrounding the nose). These artifacts limit the applicability of GE-EPI-based BOLD fMRI in pursuit of understanding the human olfactory system (Catani et al., 2013). Moreover, respiratory artifacts, primarily caused by local field shifts during inhalation and exhalation that are highly correlated with event onsets in tasked-based olfactory-related fMRI, further compromise the sensitivity of olfactory fMRI (Glover & Law, 2001).

Several techniques have been proposed to overcome these challenges. Acquiring scans at a tilted angle allows for partial compensation of local susceptibility gradients (Deichmann et al., 2003; Weiskopf et al., 2006). Alternatively, employing pulse sequences other than GE-EPI, such as T_2_-Prepared BOLD fMRI and balanced steady state free precession (bSSFP) (Parrish et al., 2008; Miao et al., 2021).

Recent work on multi-echo EPI (ME-EPI), which acquires multiple gradient echoes at each imaging timepoint, has shown potential to improve BOLD sensitivity and efficiency in resting-state fMRI (Gonzalez-Castillo et al., 2016; Lynch et al., 2020). In conventional single-echo EPI (1E-EPI), a single volume is acquired after each RF excitation at a specific echo time (TE), which determines both image signal intensity as well as BOLD contrast (Donahue et al., 2011; Gati et al., 1997; van der Zwaag et al., 2009). At a shorter TE, the image is more proton-density weighted, has less BOLD contrast, and less severe signal dropout in regions with high local susceptibility gradients. Conversely, with a longer TE, the image becomes more T_2_*-weighted, has greater BOLD contrast, and more severe signal dropout. The BOLD contrast-to-noise ratio (CNR) is expected to vary with TE, peaking at the T_2_* of each voxel (Donahue et al., 2011; Fera et al., 2004; Gati et al., 1997; Graham et al., 1996; Triantafyllou et al., 2011; van der Zwaag et al., 2009). However, various brain regions exhibit distinct T_2_* values, resulting in varying optimal TEs, making it impossible to optimize TE across the whole brain in a single-echo acquisition.

Unlike 1E-EPI, ME-EPI acquires multiple images across a range of TEs after each RF excitation, resulting in enhanced functional contrast by combining the echo images with different weights (Poser et al., 2006; Posse et al., 1999). The commonly used T_2_*-weighted echo combination method assigns the heaviest weight to the echo that is presumed to have the greatest BOLD CNR, with weighting performed separately for each voxel. By applying a T_2_*-weighted echo combination with ME-EPI, BOLD signal can be maximized at the voxel level while maintaining the temporal resolution needed for fMRI. Moreover, by leveraging the expected exponential signal decay across TEs for true BOLD contrast, noise reduction techniques can be employed to reject artifactual signals and thereby improve the BOLD contrast-to-noise ratio (DuPre et al., 2021; Gonzalez-Castillo et al., 2016; Kundu et al., 2012a, 2013, 2017; Van et al., 2023).

In this study, we implemented and tested a ME-EPI acquisition protocol and analysis pipeline specifically designed for task-based olfactory-related paradigms by incorporating shorter TE values in an effort to improve sensitivity in ventral brain regions with high static susceptibility. Using this approach in comparison to a standard 1E-EPI acquisition, we demonstrate superior sensitivity to olfactory task activation resulting in a significant sample size reduction for detecting group task activation during a representative olfactory three-alternatives forced choice task (3AFC).

## 2 Method

### 2.1 Subjects

29 subjects (13 women; aged 22 – 30 years, mean: 26, SD: 3.15) provided informed consent as approved by the University of Pennsylvania Institutional Review Board (#827217). One subject was excluded from the study due to excessive motion (number of excluded volumes, as defined in Statistical Analysis, exceeded 2% of the number of total volumes) during fMRI scanning session, leaving us with a final sample of *n* = 28. All subjects reported being right-handed nonsmokers with no history of significant neurological/psychiatric disorder or olfactory dysfunction.

### 2.2 Odor Stimuli and Delivery

Stimuli consisted of two odorants: lemon oil extract (12.3% v/v, Sigma-Aldrich, MO, USA) and benzaldehyde (0.42% v/v, Sigma-Aldrich, MO, USA). Both odorants were diluted in mineral oil (Sigma Aldrich, MO, USA). Mineral oil alone was used as a control stimulus. The odor stimuli were delivered through a custom-built computer-controlled air-diluted 12-channel olfactometer with a flow rate of 4 L/min controlled using the PsychToolbox package (Brainard, 1997) and MATLAB (Version 2020b, The Mathworks Inc., Natick, MA, USA). The medical-grade air (Airgas, Radnor, PA, USA) was first routed through two computer-controlled 6-channel gradient valve manifolds (NResearch Inc., West Caldwell, NJ, USA), regulated by a mass flow controller (MFC) (Alicat Scientific Inc., Tucson, AZ, USA). Subsequently, it passed through the headspace above 2 mL of the mineral oil-diluted odorant before being delivered to the participant through a fluorinated ethylene-propylene (FEP) plastic tube (U.S. Plastic Corp., Lima, OH, USA) placed directly below the nose.

### 2.3 Behavioral Testing Session

On Day 0, subjects participated in a behavioral testing session during which odor stimuli were presented outside the MRI environment (Figure 1a). Participants were introduced to the odors and asked to label them as well as to rate the perceived odor intensity, via the general Labeled Magnitude Scale (gLMS) (Green et al., 1993, 1996). Valence (likeability) was also assessed via the Labeled Hedonic Scale (LHS) (L. V. Jones et al., 1955; Lim et al., 2009; Schutz & Cardello, 2001; Stone et al., 2020)], as well as familiarity [Visual Analog Scale (VAS) (Dourado et al., 2021; Flint et al., 2000)]. Ratings were analyzed in MATLAB (Version 2020b, The Mathworks Inc., Natick, MA, USA), using three one-way ANOVAs to evaluate whether there were any differences among stimuli.

**Figure 1.**
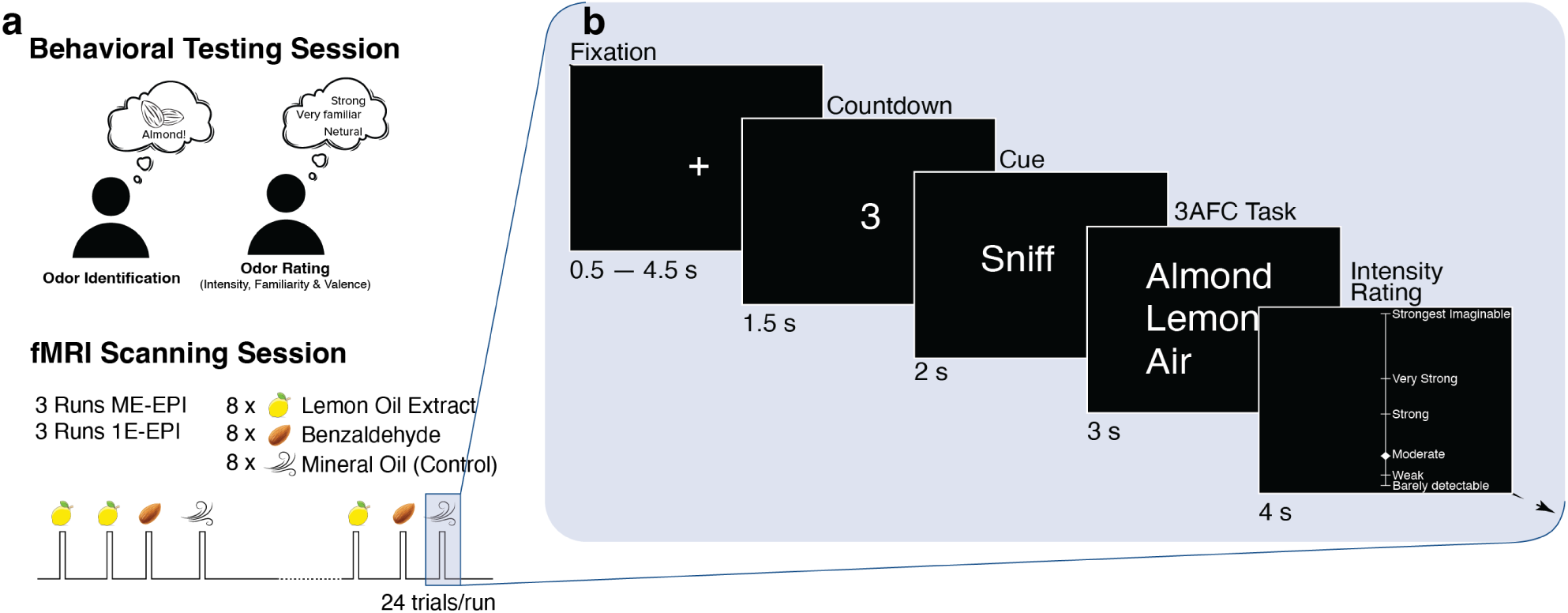
fMRI Experimental Paradigm. a) The experiment was conducted over the course of two sessions. On Day 0, the subjects participated in a series of behavioral tests where they were required to identify odors and rate their intensity, valence, and familiarity. Subsequently, an fMRI scanning session was scheduled for the subjects, taking place two to five days after their initial visit. This scanning session consisted of three runs of ME-EPI and three runs of 1E-EPI acquisition. Each run comprised 24 trials, with an equal distribution of three odor stimuli: lemon oil extract, benzaldehyde, and mineral oil. b) Timeline for a single trial. A trial started with a fixation cross presented for 0.5 – 4.5 s, followed by a 1.5 s countdown. A cue to sniff was then displayed and the odor was delivered for 2 s. The short odor onset and a long intertrial interval (11.5 s on average) were chosen to minimize habituation. Participants were then asked to identify the odor and rate its intensity.

### 2.4 fMRI Scanning Session

2-5 days after the behavioral testing session, the same participants took part in an fMRI scanning session, completing six runs, each of which contained 24 trials. On each trial, participants sniffed one of the three possible odorants (lemon oil extract, benzaldehyde, or mineral oil) and were asked to identify the odor using a three-alternative forced choice task with the labels they provided earlier during the behavioral testing session. Following the 3AFC task, they were also asked to rate the intensity of the odor. Each odor was presented eight times per run in a random order. The experimental paradigm as well as the sequence of events within a given trial is shown in Figure 1.

### 2.5 Structural and Functional MRI Data Acquisition

All magnetic resonance images were acquired using a 3 T scanner (MAGNETOM Prisma, Siemens Healthineers, Germany) using the vendor’s 64-channel head and neck coil. T_1_-weighted structural images were acquired using ME-MPRAGE (TE = 1.69/3.55/5.41/7.27 ms, TR = 2530 ms, TI = 1100 ms, FA = 8°, 1 mm isotropic).

1E-EPI and ME-EPI fMRI data were acquired using the sequence parameters outlined in Table 1. For both protocols, repetition time (TR), field of view (FOV), number of slices, resolution, flip angle (FA), echo spacing, and multiband factors are identical. To optimize the BOLD signals in OFC regions, the field of view was tilted approximately 25° to the anterior commissure-posterior commissure line (rostral > caudal) (Deichmann et al., 2003). The two sequence types were administered in an interleaved fashion in each participant, with the order being counterbalanced across participants. Of note, for 1E-EPI the echo time was set at 22 ms, which is approximately the same as the second echo time out of five echoes in the ME-EPI sequence (see Table 1). This TE aims to balance signal dropout and BOLD sensitivity for olfactory-related regions and is consistent with other olfactory-related fMRI studies (Bao et al., 2019; de Groot et al., 2021; Raithel et al., 2023; Shanahan et al., 2021; Zhou et al., 2019).

**Table 1.** fMRI Sequence Parameters. TR: Repetition time. TE: Echo time. FOV: Field of view. FA: Flip angle. 1E-EPI: single-echo EPI. ME-EPI: multi-echo EPI.

|  | 1E- EPI | ME-EPI |
| --- | --- | --- |
| **TR (ms)** | 2041 | 2041 |
| **TE (ms)** | 22 | 10.60/22.92/35.24/<br>47.56/59.88 |
| **FOV**<br>**(mm x mm)** | 216 x 216 | 216 x 216 |
| **Slices** | 48 | 48 |
| **Resolution**<br>**(mm x mm x mm)** | 2.4 x 2.4 x 2.4 | 2.4 x 2.4 x 2.4 |
| **FA (degree)** | 78 | 78 |
| **Echo Spacing**<br>**(ms)** | 0.49 | 0.49 |
| **Bandwidth**<br>**(Hz/Px)** | 2526 | 2646 |
| **Acceleration** | Multiband 2x | Multiband 2x<br>GRAPPA, 3x in-plane acceleration<br>with 36 reference lines |

Additionally, a pair of spin-echo EPI (SE-EPI) images with two phase-encode blips are acquired for both fMRI acquisition protocols (1E-EPI and ME-EPI) separately with identical FOV, number of slices, resolution, FA, echo spacing, acceleration and readout bandwidth compared to their respective fMRI acquisition. These pairs of images are then used to estimate the inhomogeneities in the static magnetic field (B_0_) during the preprocessing using a similar method described in (Andersson et al., 2003) and implemented in FSL (Smith et al., 2004) as part of fMRIPrep package (Esteban et al., 2019; Gorgolewski et al., 2011).

### 2.6 fMRI Data Preprocessing

fMRI and structural MRI data were preprocessed in each subject’s native space using fMRIPrep 21.0.0 based on Nipype 1.6.1 (Esteban et al., 2019; Gorgolewski et al., 2011). All images were then registered to ICBM 152 Nonlinear Asymmetrical template version 2009c (MNI-ICBM2009c) (2.4 mm isotropic resolution, consistent with fMRI acquisitions) before further processing (Fonov et al., 2009, 2011). Spatial smoothing was applied to all data using a Gaussian kernel of 8 mm full width at half maximum (FWHM). A detailed description of these preprocessing steps can be found in Supplementary Materials under Detailed fMRI Data Preprocessing section.

### 2.7 ME-EPI T_2_* Estimation, Echo Combination, and Multi-Echo Independent Components Analysis (ME-ICA)

Multi-echo EPI produces separate images for each echo time at each timepoint in the functional imaging run. At each timepoint, these images for the multiple echo times were combined using the T_2_*-weighted combination method (ME-WC) (Posse et al., 1999). For each voxel, T_2_* was estimated by fitting the Eq. (1) using signals from all five echoes,

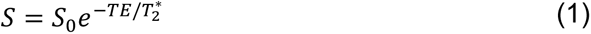

S_0_ is the initial signal magnitude without any T_2_* decay and S is the signal magnitude acquired at *t* = TE.

The images from the five echoes are then combined voxel-wise using a weighted average (ME-WC) and the weights for each echo (W_TE_) are calculated using Eq. (2) based on estimated T_2_*.

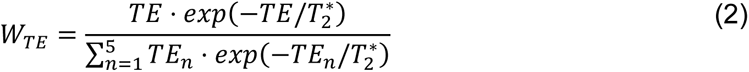

Independent components analysis (ICA) is a common strategy to decompose fMRI signals into distinct independent components, allowing the isolation of BOLD-related neuronal components from sources of noise (Beckmann, 2012; Caballero-Gaudes & Reynolds, 2017; Mckeown et al., 1998; McKeown et al., 2003). However, a persistent challenge lies in the manual classification of these independent components, which is labor-intensive and can yield inconsistent results (Kelly et al., 2010). In an attempt to tackle this problem, ME-ICA classifies each component as either a non-BOLD artifact, such as respiratory artifact, or a BOLD-like signal based on their signal characteristics across different echo times (Kundu et al., 2012b, 2013). Here, we specifically employed the TE-dependency analysis pipeline (Tedana 0.0.12) (DuPre et al., 2021) to preprocess multi-echo EPI data before registering them to the MNI template.

In the original TEDANA pipeline, a monoexponential model [Eq. (1)] was fit to the data at each voxel using nonlinear model fitting to estimate T_2_* and S_0_ maps, using T_2_*/S_0_ estimates from a log-linear fit as initial values. This approach generally performs well across most brain regions. However, in regions with significant signal dropout at longer echo times, particularly due to susceptibility artifacts in areas such as the orbitofrontal cortex and entorhinal cortex, this fitting method tends to overestimate T_2_*.

Consequently, we replaced the TEDANA estimate of T2* with our own estimates, generated by fitting the following model voxel-wise:

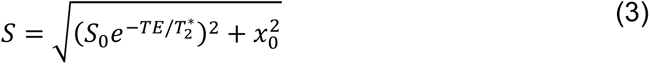

In this model, *x_0_* represents the thermal noise floor, which does not decay over TE and whose contribution to the overall signal increases in later echoes as the signal diminishes in high-susceptibility regions. This model was fit to the data by minimizing the 2-norm of the weighted residuals [Eq. (4)] via the Trust-Region-Reflective algorithm (Coleman & Li, 1996).

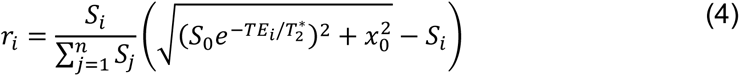

The weighted residuals are calculated as the difference between the estimated signals *Ŝ_i_* and the observed signals *S_i_*, weighted by the observed signals *S_i_* across all *n* = 1000 observations (200 volumes/run, 5 echoes/volume).

### 2.8 Estimating Motion for Different Odor Stimuli

The framewise displacement (FD), calculated during fMRI preprocessing using the method described in (Power et al., 2012), and the onset of odor stimulus for each condition were extracted for each run. The coefficient of determination (R^2^) was then estimated by calculating the Pearson correlation coefficient between the FD and the odor onset for each stimulus condition convolved with canonical hemodynamic response function (HRF) for each subject (Pearson, 1895; Siegel, 1956). To compare whether there is a group-level difference in movement parameters for different odor stimuli, two-sided paired *t*-tests were performed on the R^2^, across all three stimulus conditions ([Lemon vs. Control], [Benzaldehyde vs. Control], and [Lemon vs. Benzaldehyde]) for both 1E-EPI and ME-EPI acquisitions.

### 2.9 Physiological Data Analysis

The respiratory trace was recorded with a respiratory transducer using a pressure sensor (BIOPAC System, CA, USA) affixed around the subject’s chest, and then preprocessed before being added as a nuisance regressor: the trace was smoothed using a moving average with a 250 ms window, high-pass filtered at 0.05 Hz with an infinite impulse response (IIR) filter, detrended, and standardized such that the mean of the signals is 0 and the standard deviation is 1 (Liu et al., 2024; Raithel et al., 2023). The preprocessed trace was also downsampled to 1/2.4 Hz to match the TR of the fMRI acquisitions and included as a nuisance regressor. Two additional nuisance regressors were introduced: the trial-by-trial sniff volume was defined as the amplitude difference between the peak and the trough within the cued sniff, and the sniff duration was defined as the time difference between the peak and the trough.

### 2.10 Statistical Analysis

All fMRI data were analyzed in SPM12 using a general linear model (GLM) to estimate the main effects of the odor stimulation. Each experimental condition (control, lemon oil, and benzaldehyde) was modeled using an independent regressor for each run. This results in a 2 (multi-echo and single-echo) by 3 (control, lemon oil, and benzaldehyde) factorial design. The model included the following nuisance regressors: trial-by-trial sniff volume and sniff duration, both convolved with the hemodynamic response function (HRF), as well as the down-sampled breathing trace and 24 movement parameters (six movement parameters derived from spatial realignment, their squares, derivatives, and squares of derivatives). Additional nuisance regressors were introduced to exclude volumes with excessive motion, defined as those with at least one of six movement parameters derived from spatial realignment deviating by more than 6 standard deviations from the mean. In all GLMs, data were high-pass filtered at 1/128 Hz; the temporal autocorrelation was estimated and removed as an autoregressive AR(1) process globally (Friston et al., 2002).

Contrasts of interest were defined as [Odor > Control], [Lemon > Control] and [Benzaldehyde > Control]. Whole brain analysis was performed to validate that odor-evoked activations were present in the data acquired through both protocols (1E-EPI and ME-EPI). Small volume corrections (SVC) and ROI analysis were separately performed on OFC, amygdala, piriform cortex and entorhinal cortex, as suggested by previous studies highlighting the involvement of these regions in human olfactory processing (Anderson et al., 2003). Amygdala and piriform cortex were defined anatomically using a human brain atlas (Mai et al., 2016). Delineation of OFC and entorhinal cortex were based on the MNI-ICBM2009c atlas (Manera et al., 2020).

### 2.11 Post-hoc Power Analysis

We used our ROI analysis results to estimate the sample sizes required to detect significant olfactory-related group fMRI activation in subsequent studies. A priori power analyses were performed with α = 0.05 and β = 0.80 using G*Power (Faul et al., 2009) for both ME-EPI and 1E-EPI data in the same ROIs described in the previous section.

## 3 Results

### 3.1 Echo-combined Images Reduce Signal Dropout

We first examined the estimated T_2_* values for each olfactory-related region. As expected, signal dropouts are reduced in shorter echo times (Figure 2a, TE_1_ and TE_2_), while T_2_*-weighted contrast is greater in later echo times (Figure 2a, TE_3_ – TE_5_). In particular, the medial and lateral OFC along with the entorhinal cortex, exhibited notably smaller T_2_* values due to susceptibility artifacts (Figure 2a & 2b), and in turn the echo-combined image more heavily weights the shorter echoes (Figure 2c). Using these weights, the combined image (Figure 2a, ME-WC) effectively mitigated signal dropout while retaining notable T_2_*-weighted contrast.

**Figure 2.**
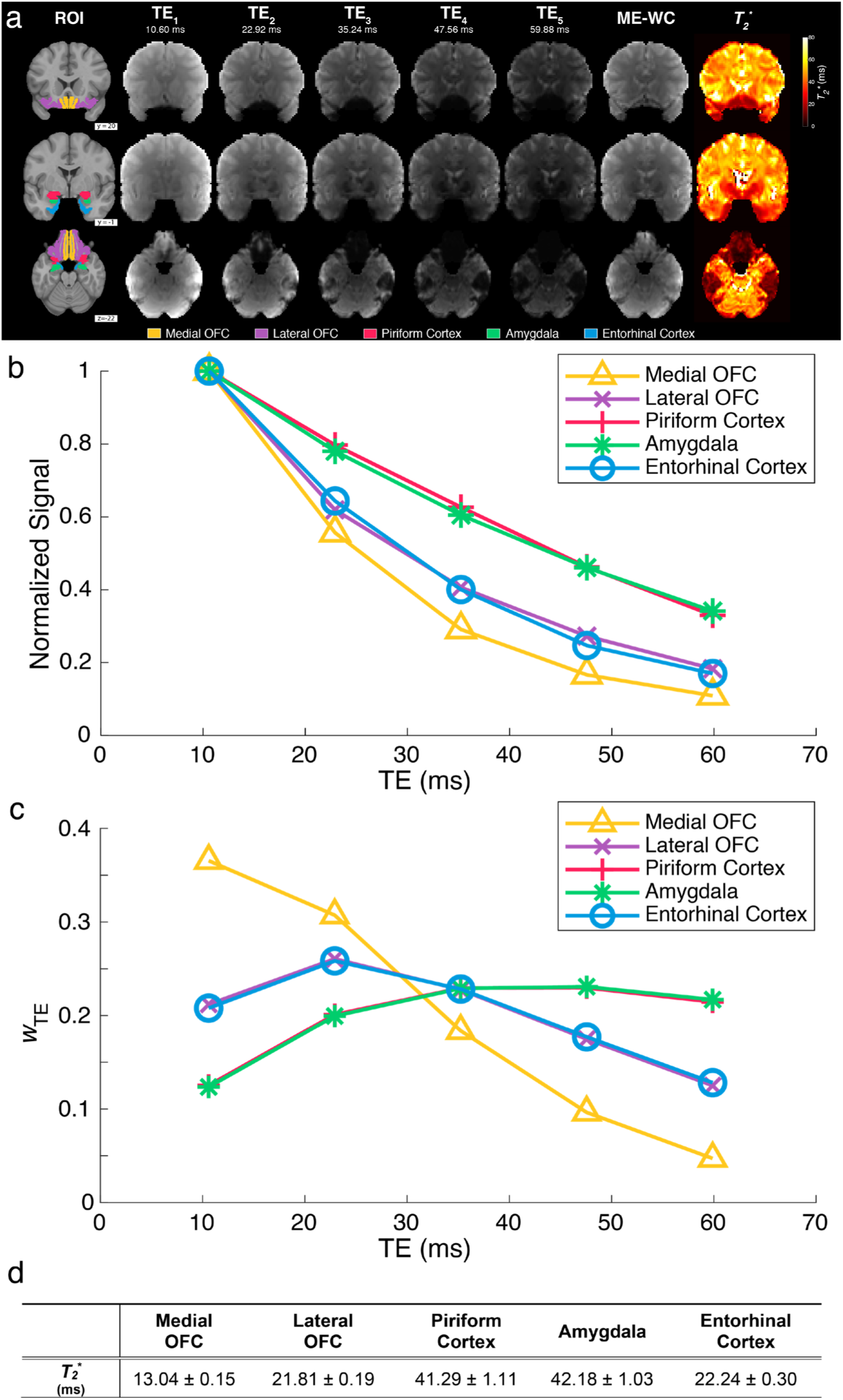
Multi-Echo fMRI Reduces Signal Dropout and Improves BOLD Contrast through T_2_*-Weighted Combination in a Single Subject. a) Two representative coronal slices containing the regions of interest are shown with a variety of contrasts: ROIs overlayed on the T_1_-weighted image (ROI); estimated T_2_* maps; all five individual echo images (TE_1_ – TE_5_) from the ME-EPI acquisition; and the T_2_*-weighted combined images (ME-WC). b) Plot of signal vs echo time for each anatomical ROI within a single subject. Within each ROI, mean signals for each individual echo were normalized by dividing by the mean signal from the first echo. c) Within-ROI mean of the voxel-wise weights used to compute the T_2_ *-weighted combined images for the same subject. d) Estimated mean T_2_* ± 95% confidence interval for each ROI.

### 3.2 BOLD Activation in Diverse Olfactory Brain Regions During Odor Stimulation

We next investigated the regions that were significantly activated when the odors were delivered under ME-EPI acquisition with the ME-ICA denoising method [Odor > Control] (Figure 3a) (DuPre et al., 2021). Consistent with previous literature, we observed significant activation in the piriform cortex, amygdala, entorhinal cortex, insula, and OFC bilaterally (*p* < 0.001, uncorrected) (Anderson et al., 2003; Dikecligil & Gottfried, 2024; Gottfried et al., 2003; Howard & Gottfried, 2014; Patin & Pause, 2015; Plailly et al., 2008). We next examined the BOLD effects of individual odor stimulus, namely [Benzaldehyde > Control] and [Lemon > Control]. Despite statistical tests indicating no significant differences in perceived intensity (*p* = 0.38), valence (*p* = 0.45), and familiarity (*p* = 0.11) between benzaldehyde and lemon oil and no significant differences in motion among three experimental conditions ([Lemon vs. Control] *p* = 0.60, [Benzaldehyde vs. Control] *p* = 0.33, and [Lemon vs. Benzaldehyde] *p* = 0.63), our analysis revealed that the main effect [Odor > Control] was driven mainly by the contrast [Lemon > Control]. In comparison, the contrast [Benzaldehyde > Control] produced significant levels of activation only in the bilateral insular region (Figure 3a).

**Figure 3.**
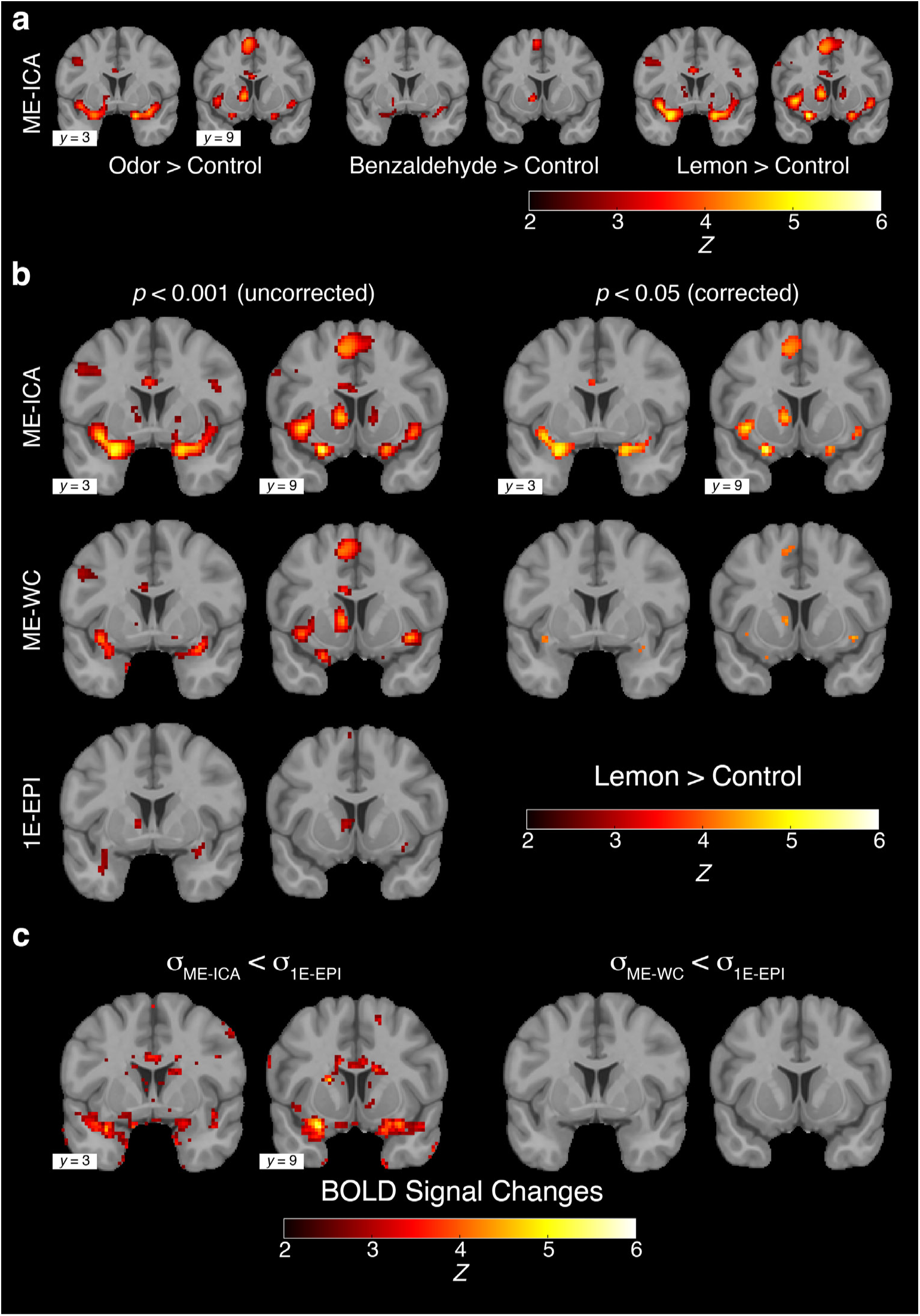
Activation Maps from Olfactory-Related Task for Both 1E-EPI and ME-EPI. a) Two representative coronal slices of activation maps for all three contrasts ([Odor > Control], [Benzaldehyde > Control], [Lemon > Control]) using ME-EPI acquisition with ME-ICA denoising methods (*p* < 0.05, FDR corrected). b) Activation maps for [Lemon > Control] contrast for ME-EPI acquisition with ME-ICA denoising methods (ME-ICA), and without ME-ICA denoising methods (ME-WC) as well as single-echo fMRI acquisition (1E-EPI). The left two columns show the maps without multiple comparison correction (*p* < 0.001) while the right two columns show the maps corrected for multiple comparison (*p* < 0.05, FDR corrected). No activation survived correction for multiple comparisons for 1E-EPI (not shown). c) Z-score statistical maps yielded from Levene’s test to examine whether BOLD signal variance was lower in ME-WC [σ_ME-WC_ < σ_1E-EPI_] and ME-EPI [σ_ME-ICA_ < σ_1E-EPI_] compared to 1E-EPI (*p* < 0.05, FDR corrected).

The difference in BOLD response to these stimuli was not anticipated though not wholly unprecedented, as previous research has demonstrated that different odors can elicit distinct responses within olfactory brain regions even in the absence of differences in low-level stimulus features (Fournel et al., 2016; Howard & Gottfried, 2014; Savic, 2002; Savic et al., 2002). In the single-echo fMRI, a similar trend is observed, with more active regions being detected with lemon oil (*p* < 0.05, uncorrected, Figure S2). To our knowledge, there is no literature in humans that directly elicits piriform cortex BOLD activation with a similar sample size for benzaldehyde. However, rodent studies have reported that benzaldehyde demonstrates BOLD activation in the amygdala, as well as the piriform and entorhinal cortices (Kulkarni et al., 2012). Accordingly, to compare and observe the differences between acquisition methods within olfactory regions, we will focus on the activation under lemon oil condition [Lemon > Control] for our further analyses.

### 3.3 ME-EPI Improves fMRI Signal Detection by Both Increasing BOLD Signal Sensitivity and Specificity

Having validated that our ME-EPI acquisition protocol is suitable to capture odor-evoked BOLD responses, we next compared BOLD activation among different acquisition methods (1E-EPI vs. ME-EPI) and analysis pipelines (ME-WC vs. ME-ICA). A whole-brain analysis using either whole-brain multiple comparisons correction with false discovery rate (FDR) (Benjamini & Hochberg, 1995; Benjamini & Yekutieli, 2001; Yekutieli & Benjamini, 1999) or small volume correction (SVC) did not yield any supra-threshold voxels for 1E-EPI. However, the same whole-brain analysis using the T_2_*-weighted combined ME-EPI (ME-WC) highlighted activation in all ROIs (piriform cortex, amygdala, OFC, and entorhinal cortex) (Figure 3b). Moreover, by using ME-ICA denoising methods (DuPre et al., 2021), we observed a further increase in both the number of activation voxels and peak Z values in all ROIs (Table 2).

**Table 2.** Number of Voxels and Peak Z-Score Activated by Lemon Oil for each ROI (p < 0.05, SVC).

|  | Piriform Cortex |  | Amygdala |  | Entorhinal Cortex |  | OFC |  |
| --- | --- | --- | --- | --- | --- | --- | --- | --- |
|  | # Voxel | Peak Z | # Voxel | Peak Z | # Voxel | Peak Z | # Voxel | Peak Z |
| **ME-ICA** | 158 | 5.43 | 6 | 4.21 | 11 | 4.78 | 27 | 5.43 |
| **ME-WC** | 4 | 4.32 | 2 | 3.63 | 1 | 3.82 | 20 | 4.62 |
| **1E-EPI** | 0 | N/A | 0 | N/A | 0 | N/A | 0 | N/A |

We next examined potential reasons for the observed differences. Specifically, we hypothesized the improvement of the BOLD sensitivity in the case of the ME-EPI data was due to larger BOLD signal changes (i.e., greater sensitivity), as we performed a locally weighted combination of all echoes for each individual voxel instead of using one single TE across the brain. BOLD signal changes could be calculated by contrast estimates from the GLM. Therefore, we performed a paired *t*-test on contrast estimates [Lemon > Control] between 1E-and ME-EPI with ME-ICA [ME-ICA > 1E-EPI] among all ROIs. Only piriform cortex (*p* = 0.023) and lateral OFC (*p* = 0.006) showed greater estimates for the contrast [Lemon > Control] for ME-ICA compared to 1E-EPI.

An alternative explanation for the observed increase in BOLD detection for ME-EPI versus 1E-EPI, may lie in the reduction of the background noise due to non-BOLD signals (i.e., greater specificity). To test the hypothesis that ME-EPI signals have lower variance, we performed a Levene’s test (Levene, 1960). This test is commonly employed to assess whether two or more groups have equal variances in a dataset (Chiarello et al., 2009; Farinha et al., 2022; Im et al., 2010; T. B. Jones et al., 2010; Swarup et al., 2013). When using a whole-brain analysis to compare the signal variance between ME-ICA and 1E-EPI, we observed suprathreshold clusters (*p* < 0.05, FDR corrected) overlapping with the voxels activated for the [Lemon > Control] contrast (*p* < 0.05, FDR corrected) from ME-ICA (Figure 3c). We then generated a mask using those activated voxels and found that 18.89% of the masked voxels reached the significance threshold under Levene’s test. This suggests that ME-ICA decreases the signal variance, thereby reducing non-BOLD noise and increasing BOLD specificity. We did not observe any suprathreshold clusters when comparing ME-WC and 1E-EPI.

Taken together, these tests suggest that our observed increase in BOLD detection is not only due to increased sensitivity BOLD signals, but also the reduction of non-BOLD artifacts enhancing BOLD specificity.

### 3.4 ME-EPI Improves Statistical Power and Reduces Sample Sizes

So far, we have established that ME-EPI improved BOLD signal sensitivity by both enhancing BOLD signal and reducing non-BOLD artifacts. An outstanding question is to what degree BOLD sensitivity is improved when using our suggested method in the context of task-based olfactory fMRI. To quantify the change in BOLD sensitivity, we examined the contrast [Lemon > Control] in the brain regions central to olfaction, namely the piriform cortex, amygdala, lateral and medial OFC, and entorhinal cortex for both acquisition methods separately and then compared the statistical outcomes. Whereas activation in medial OFC did not reach statistical significance using data resulting from either acquisition method, all remaining regions show much lower *p*-values for ME-ICA compared to 1E-EPI (Table 3). Furthermore, to determine the minimum sample size required to observe significance in each region, post-hoc power analyses were carried out (α = 0.05 and β = 0.80) to estimate the effect of each acquisition method on the sample sizes needed to detect effects in similar olfactory task fMRI studies. ME-ICA led to a substantial reduction in the sample size, ranging from by half to one-fourth, compared to 1E-EPI, depending on the specific regions (Figure 4).

**Figure 4.**
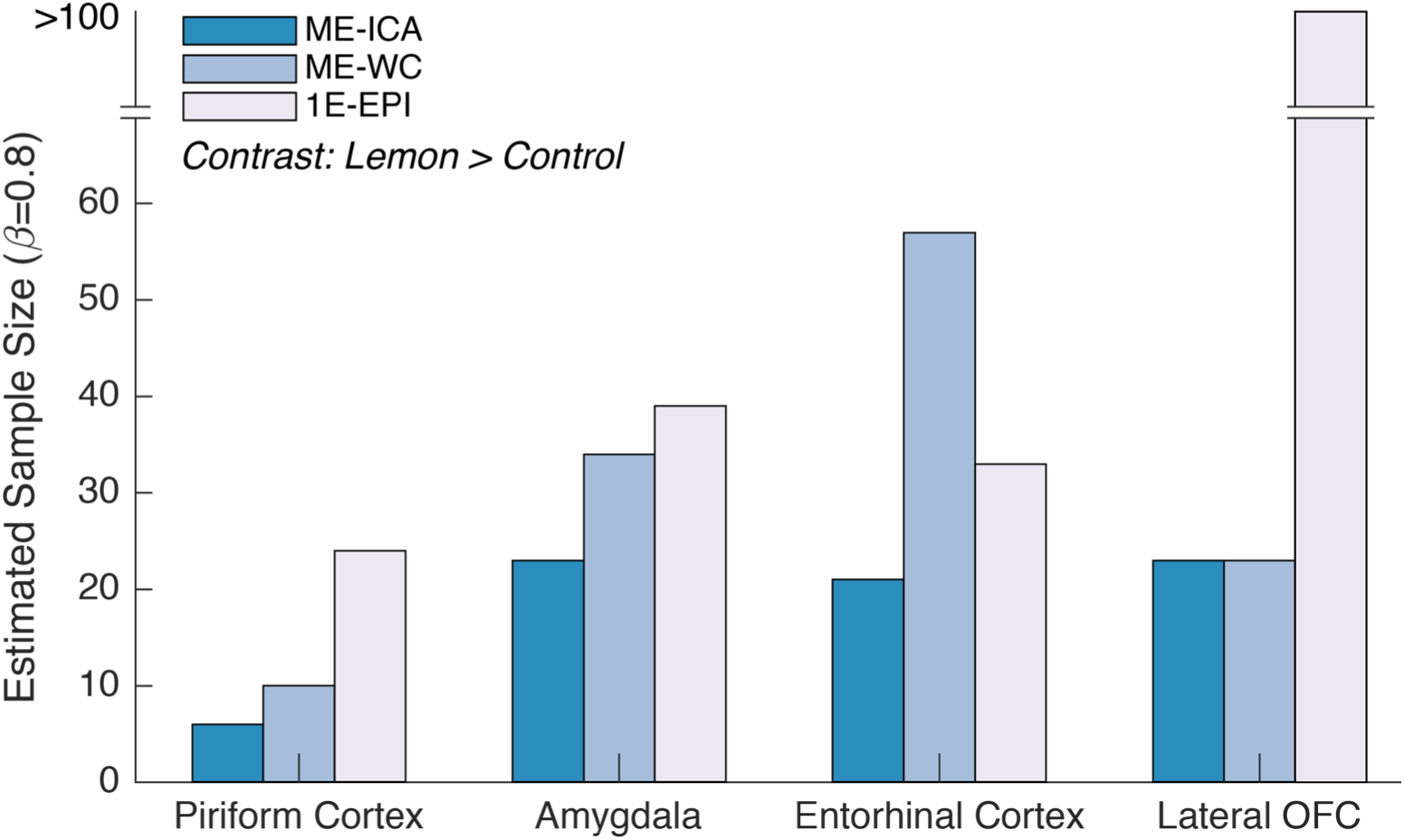
Estimated Sample Size for ROI Analyses Based on Post-hoc Power Analyses (α = 0.05 and β = 0.80) in all Olfactory-Related ROIs under Contrast [Lemon > Control] for Single-echo fMRI (1E-EPI), ME-EPI BOLD Acquisition with ME-ICA Denoising Methods (ME-ICA), and without ME-ICA Denoising Methods (ME-WC). The estimated sample sizes for medial OFC for 1E-EPI, ME-ICA and ME-WC were not shown since they were all above 100.

**Table 3.** Statistical Significance of ROI Analyses for Each Olfactory-related ROI under the Contrast [Lemon > Control] for 1E-EPI, ME-WC and ME-ICA.

|  | Piriform Cortex | Amygdala | Entorhinal Cortex | Lateral OFC | Medial OFC |
| --- | --- | --- | --- | --- | --- |
| **ME-ICA** | < 0.0001*** | 0.0052** | 0.0039** | 0.0058** | 0.7663 |
| **ME-WC** | 0.0001*** | 0.0173* | 0.0505 | 0.0054** | 0.5607 |
| **1E-EPI** | 0.0066** | 0.0243* | 0.0157* | 0.5429 | 0.6687 |

## 4 Discussion

Susceptibility artifacts pose a significant challenge when using GE-EPI for fMRI studies. The primary olfactory cortex not only suffers from some of the most severe static susceptibility artifacts within the human brain but is also affected by respiration-related artifacts. Here, we have developed and validated a ME-EPI protocol and analysis pipeline specifically tailored for olfactory task-based fMRI. In the context of a basic olfactory 3AFC task, we found that ME-EPI, alongside ME-ICA denoising, systemically improves BOLD sensitivity and reduces the required sample sizes needed to infer a significant effect at a group level compared to conventional 1E-EPI. This is achieved by both increasing BOLD signal sensitivity and reducing the contributions of non-BOLD signal changes using the additional denoising features ME-EPI provided.

The echo times used for ME-BOLD provided a range of contrasts across different TEs for each ROI of interest (Figure 2a). When evaluating the T_2_*-derived weights calculated for the echo-combined image, we confirmed that high-susceptibility regions with short T_2_* values, namely, lateral and medial OFC, and entorhinal cortex, had higher weights assigned to the first two echoes. In contrast, the remaining ROIs had higher weights assigned to the remaining (and later) three echoes. As a result, the T_2_*-weighted combined (ME-WC) images were able to produce more uniform T_2_*-weighted contrast across the whole brain (Figure 2a, ME-WC).

When investigating the region activated by odor onset using ME-ICA, we detected activation in all major primary olfactory cortex regions (piriform cortex, amygdala, entorhinal cortex, and OFC bilaterally) in addition to the insular region (Figure 3a). These findings are consistent with previous literature (Anderson et al., 2003; Dikecligil & Gottfried, 2024; Gottfried et al., 2003; Howard & Gottfried, 2014; Patin & Pause, 2015; Plailly et al., 2008). Further analysis revealed that the activation was primarily driven by the lemon extract stimulus in comparison to benzaldehyde. Previous studies have demonstrated that different odor stimuli can activate distinct primary olfactory regions to different degrees (Fournel et al., 2016; Howard & Gottfried, 2014; Savic, 2002; Savic et al., 2002). In this study we found that that benzaldehyde activates the olfactory system to a lesser extent than lemon oil, such that the observed activation did not reach significance after correcting for multiple comparisons in the whole brain analysis.

When comparing the results of 1E-EPI and ME-EPI, we observed significantly increased sensitivity in both whole brain analysis and ROI analysis, which are two commonly used univariate fMRI analysis methods. In the case of whole brain analysis (Figure 3b), we found that ME-EPI exhibited higher peak Z values and a greater number of activated voxels compared to 1E-EPI, as shown in Table 2. It is worth noting that even without using the ME-ICA denoising method, ME-EPI still outperformed 1E-EPI by performing a T_2_*-weighted combination of data from all echoes. This improvement was expected, as the weighting of data from each echo time was designed to enhance the sensitivity to BOLD contrast (Posse et al., 1999).

For ROI analysis and sequential power analysis, which were used to estimate minimal sample sizes, ME-EPI with the ICA denoising method demonstrated superiority over 1E-EPI, as shown in Table 3 and Figure 4. We attribute these improvements to two factors: the increase of BOLD signal intensity through T_2_*-weighted echo combination and the reduction of noise by leveraging signal evaluation across the echo times. To support these hypotheses, we conducted a series of statistical tests (Figure 3c) to show the mean signals are significantly higher and the variance is significantly lower in ME-ICA acquisition compared to 1E-EPI.

While both ME-EPI and ME-WC overwhelmingly increased the statistical power for ROI analysis, we observed an unexpected decrease in statistical power for ME-WC in the entorhinal cortex. Previous studies have demonstrated that local B_0_ field shifts affect T_2_* values near the ethmoid sinuses during respiration, resulting in fluctuations of T_2_* values across respiratory cycles (Barry & Menon, 2005; Raj et al., 2000, 2001; Van de Moortele et al., 2002). The T_2_*-weighted combination method used in our analyses adopted a fixed weighting approach throughout a single scan, potentially failing to account for these dynamic T_2_* fluctuations. However, our additional analyses showed that, although statistically significant in most ROIs, T_2_* values exhibit only subtle differences between inhalation and exhalation phases (Table S1). This, in return, demonstrates the robustness of the echo combination method against respiratory phase variations.

We also evaluated whether nonexponential signal decay contributed to suboptimal echo combination weightings, leading to the unexpected result. As shown in Figure S2, signals from most regions fit well with exponential decay, but regions with high susceptibility, such as the entorhinal cortex, medial, and lateral OFC, exhibited more pronounced nonexponential decay. Yet, this observation does not fully explain the unexpected decreases observed exclusively in the entorhinal cortex.

Furthermore, we examined the effects of ME-WC and ME-ICA on the distribution of statistical significance within each ROI (Figure S1). We have shown that ME-WC increases sensitivity to both positively correlated BOLD activation and anti-correlated signals. Moreover, ME-WC and ME-ICA both increases the peak Z-scores and the number of statistically significant voxels, consistent with voxel-wise whole-brain analysis (Table 2 and Figure 3b).

Additionally, we investigated the effect of the number of echoes on the main effect, considering whether later echoes contribute significantly. Using the same analysis pipeline, we compared whole-brain analysis results from data including the first 3, 4, and all 5 echoes (Figure S4). As the number of echoes increased, the size of the clusters also increased.

The current analysis pipeline utilized a specific method of combining multiple echo image series into a single image series: T_2_*-weighted combination, also known as “optimal combination”, is used in most ME-EPI studies as it is designed to maximize BOLD-contrast sensitivity (Baracchini et al., 2021; Gonzalez-Castillo et al., 2016; Kundu et al., 2017; Lynch et al., 2020; Roth et al., 2020). However, an interesting finding emerged when we processed the ME-EPI data separately and treated each individual echo series as a 1E-EPI dataset. Surprisingly, no supra-threshold activation was detected at the first echo (*p* < 0.05, uncorrected), while the majority of activations were observed at the later echoes (*p* < 0.05, uncorrected) (Figure S5). When examining the Z values for the ROI analyses across all 5 individual echoes, we also observe a similar trend (Table S2). However, regions with short T_2_* (such as the lateral and medial OFC and entorhinal cortex in this study) predominantly assigned higher weights to the first two echoes.

Interestingly, in the process of understanding the T_2_*-weighted combination method implemented in the TEDANA package, we observed that directly estimating T_2_* by fitting the Eq. (1) tends to overestimate T_2_* in regions with high susceptibility (Table S3). As described in the Method, we proposed using Eq. (3) as the objective function to estimate T_2_* for each voxel. While this approach improved some estimations, work is needed to improve the accuracy of these estimations in the future.

There have also been recent advances in denoising methods that leverage phase information, such as Noise Reduction with Distribution Corrected (NORDIC) PCA (Dowdle et al., 2023; Moeller et al., 2021; Vizioli et al., 2021). These methods have demonstrated the ability to further enhance BOLD sensitivity in resting-state fMRI. However, our study did not acquire phase data, which prevented us from testing these advanced denoising methods in our data.

In this study, we utilized a limited set of two odor stimuli, along with a control stimulus. Although this approach was adequate to compare the ME-EPI and 1E-EPI acquisition methods, future studies will aim to incorporate a wider range of odor stimuli. By including multiple odor stimuli, we can expand our analysis to encompass multivariate analysis, such as Multivariate Pattern Analysis (MVPA), which has been commonly utilized in other olfaction task-based fMRI studies (Donoshita et al., 2021; Howard et al., 2009; Howard & Gottfried, 2014; Perszyk et al., 2023), though we expect also to see an improvement in multivariate analysis using ME-ICA.

In conclusion, our study demonstrated the utility of a ME-EPI protocol combined with a pre-configured analysis pipeline for fMRI in olfactory cortex. This approach enhanced sensitivity to task-dependent BOLD contrast changes in olfactory regions, and post-hoc power analyses indicate that group effects can be detected with a reduced sample size compared to standard 1E-EPI. Although our protocol and pipeline were specifically designed for task-based fMRI in the presence of large static susceptibility, a longstanding challenge in the field of fMRI, they showcase the superior denoising capabilities of ME-EPI for task-based fMRI more generally. These findings strongly support the adoption of ME-EPI over 1E-EPI for future task-based fMRI studies.

## Data and Code Availability

The unthresholded Z-maps showing the whole-brain results for univariate analyses were made available on Neurovault (https://neurovault.org/collections/KGZMOMWR/). The modified TEDANA codes with updated T2* estimation methods described in Section 2.7 are available on GitHub (https://github.com/luxwig/tedana/tree/altT2s).

## Author Contributions

**Ludwig Sichen Zhao**: Conceptualization, Methodology, Software, Formal Analysis, Investigation, Data Curation, Visualization, Writing – Original Draft, Project administration. **Clara U. Raithel**: Conceptualization, Methodology, Investigation, Writing – Review & Editing. **M. Dylan Tisdall**: Conceptualization, Methodology, Writing – Review & Editing, Supervision, Resources. **John A. Detre**: Conceptualization, Methodology, Writing – Review & Editing, Supervision, Resources. **Jay A. Gottfried**: Conceptualization, Methodology, Writing – Review & Editing, Supervision, Resources, Funding acquisition

## 1 Detailed fMRI Data Preprocessing

Results included in this manuscript come from preprocessing performed using *fMRIPrep* 21.0.0 (Esteban, Markiewicz, et al. (2018); Esteban, Blair, et al. (2018); RRID:SCR_016216), which is based on *Nipype* 1.6.1 (K. Gorgolewski et al. (2011); K. J. Gorgolewski et al. (2018); RRID:SCR_002502).

### 1.1 Preprocessing of B_0_ Inhomogeneity Mappings

A total of 2 fieldmaps were found available within the input BIDS structure for this particular subject. A B_0_-nonuniformity map (or *fieldmap*) was estimated based on two (or more) echo-planar imaging (EPI) references with topup (Andersson, Skare, and Ashburner (2003); FSL 6.0.5.1:57b01774).

### 1.2 Anatomical Data Preprocessing

A total of 1 T1-weighted (T1w) images were found within the input BIDS dataset. The T1-weighted (T1w) image was corrected for intensity non-uniformity (INU) with N4BiasFieldCorrection (Tustison et al. 2010), distributed with ANTs 2.3.3 (Avants et al. 2008, RRID:SCR_004757), and used as T1w-reference throughout the workflow. The T1w-reference was then skull-stripped with a *Nipype* implementation of the antsBrainExtraction.sh workflow (from ANTs), using OASIS30ANTs as target template. Brain tissue segmentation of cerebrospinal fluid (CSF), white-matter (WM) and gray-matter (GM) was performed on the brain-extracted T1w using fast (FSL 6.0.5.1:57b01774, RRID:SCR_002823, Zhang, Brady, and Smith 2001). Brain surfaces were reconstructed using recon-all (FreeSurfer 6.0.1, RRID:SCR_001847, Dale, Fischl, and Sereno 1999), and the brain mask estimated previously was refined with a custom variation of the method to reconcile ANTs-derived and FreeSurfer-derived segmentations of the cortical gray-matter of Mindboggle (RRID:SCR_002438, Klein et al. 2017). Volume-based spatial normalization to one standard space (MNI152NLin2009cAsym) was performed through nonlinear registration with antsRegistration (ANTs 2.3.3), using brain-extracted versions of both T1w reference and the T1w template. The following template was selected for spatial normalization: *ICBM 152 Nonlinear Asymmetrical template version 2009c* [Fonov et al. (2009), RRID:SCR_008796; TemplateFlow ID: MNI152NLin2009cAsym].

### 1.3 Functional Data Preprocessing

For each of the 6 BOLD runs found per subject (across all tasks and sessions), the following preprocessing was performed. First, a reference volume and its skull-stripped version were generated by aligning and averaging 5 single-band references (SBRefs). For single-echo dataset, the reference volume is the single-band references (SBRefs). Head-motion parameters with respect to the BOLD reference (transformation matrices, and six corresponding rotation and translation parameters) are estimated before any spatiotemporal filtering using mcflirt (FSL 6.0.5.1:57b01774, Jenkinson et al. 2002). The estimated *fieldmap* was then aligned with rigid-registration to the target EPI (echo-planar imaging) reference run. The field coefficients were mapped on to the reference EPI using the transform. BOLD runs were slice-time corrected to 0.969s (0.5 of slice acquisition range 0s-1.94s) using 3dTshift from AFNI (Cox and Hyde 1997, RRID:SCR_005927). The BOLD reference was then co-registered to the T1w reference using bbregister (FreeSurfer) which implements boundary-based registration (Greve and Fischl 2009). Co-registration was configured with six degrees of freedom. First, a reference volume and its skull-stripped version were generated using a custom methodology of *fMRIPrep*. Several confounding time-series were calculated based on the *preprocessed BOLD*: framewise displacement (FD), DVARS and three region-wise global signals. FD was computed using two formulations following Power (absolute sum of relative motions, Power et al. (2014)) and Jenkinson (relative root mean square displacement between affines, Jenkinson et al. (2002)). FD and DVARS are calculated for each functional run, both using their implementations in *Nipype* (following the definitions by Power et al. 2014). The three global signals are extracted within the CSF, the WM, and the whole-brain masks. Additionally, a set of physiological regressors were extracted to allow for component-based noise correction (*CompCor*, Behzadi et al. 2007). Principal components are estimated after high-pass filtering the *preprocessed BOLD* time-series (using a discrete cosine filter with 128s cut-off) for the two *CompCor* variants: temporal (tCompCor) and anatomical (aCompCor). tCompCor components are then calculated from the top 2% variable voxels within the brain mask. For aCompCor, three probabilistic masks (CSF, WM and combined CSF+WM) are generated in anatomical space. The implementation differs from that of Behzadi et al. in that instead of eroding the masks by 2 pixels on BOLD space, the aCompCor masks are subtracted a mask of pixels that likely contain a volume fraction of GM. This mask is obtained by dilating a GM mask extracted from the FreeSurfer’s *aseg* segmentation, and it ensures components are not extracted from voxels containing a minimal fraction of GM. Finally, these masks are resampled into BOLD space and binarized by thresholding at 0.99 (as in the original implementation). Components are also calculated separately within the WM and CSF masks. For each CompCor decomposition, the *k* components with the largest singular values are retained, such that the retained components’ time series are sufficient to explain 50 percent of variance across the nuisance mask (CSF, WM, combined, or temporal). The remaining components are dropped from consideration. The head-motion estimates calculated in the correction step were also placed within the corresponding confounds file. The confound time series derived from head motion estimates and global signals were expanded with the inclusion of temporal derivatives and quadratic terms for each (Satterthwaite et al. 2013). Frames that exceeded a threshold of 0.5 mm FD or 1.5 standardised DVARS were annotated as motion outliers. All resamplings can be performed with *a single interpolation step* by composing all the pertinent transformations (i.e. head-motion transform matrices, susceptibility distortion correction when available, and co-registrations to anatomical and output spaces). Gridded (volumetric) resamplings were performed using antsApplyTransforms (ANTs), configured with Lanczos interpolation to minimize the smoothing effects of other kernels (Lanczos 1964). Non-gridded (surface) resamplings were performed using mri_vol2surf (FreeSurfer).

Many internal operations of *fMRIPrep* use *Nilearn* 0.8.1 (Abraham et al. 2014, RRID:SCR_001362), mostly within the functional processing workflow. For more details of the pipeline, see the section corresponding to workflows in *fMRIPrep*’s documentation.

## 2 Impacts of ME-WC and ME-ICA on the Distribution of Statistical Significance

In Figure S1, the number of voxels (cluster size) exceeding the statistical significance thresholds (indicated by the dashed lines) and the peak Z-scores are higher for ME-WC compared to 1E-EPI. ME-ICA further increases the number of voxels above these thresholds, which is consistent with the results from voxel-wise whole brain analysis (Table 2 and Figure 3b). When examining the mean values and the 95% confidence intervals (CI) of the distribution, we observed that ME-WC has more voxels shifting left past the thresholds, increasing the suprathreshold cluster size as well as peak Z-scores and thus widening the distribution and increasing variance. In other words, ME-WC tends to have a wider CI, indicating larger variance and a broader spread of the distribution.

The significance of the ROI analysis depends on both the mean and the variance of the distribution. In some regions, such as the entorhinal cortex, ME-WC increases the within-ROI variance without increasing the mean, and thus the statistical significance decreases compared to 1E-EPI. We interpret this as meaning that while ME-WC has increased sensitivity to positively correlated BOLD activation, it has also increased the sensitivity to anti-correlated signals.

## 3 T_2_* Changes Due to Respiration

To investigate T_2_ * changes during respiration, we estimated T_2_* maps for each subject at both inhalation and exhalation phases. As described in the Methods section, the respiratory trace was downsampled to match the TR of the fMRI acquisition. Inhalation phase volumes are defined as those 0.8 standard deviations above the mean of the downsampled respiratory trace, while exhalation phase volumes are defined as those 0.8 standard deviations below the mean.

We extracted the mean T_2_* values for each subject at both inhalation and exhalation phases and performed two-sample, two-sided *t*-tests to determine if there were statistically significant differences between the two phases. All ROIs, except the lateral orbitofrontal cortices (OFC), showed significant, though subtle, differences, with increased T_2_* values during the inhalation phase, as shown in Table S1. As a result, the weights in the T_2_*-weighted combination method (ME-WC) remain relatively stable across different respiratory phases.

## 4 Nonexponential Decay of Multi-Echo GE-EPI Signal

Theoretically, the BOLD signal under GE-EPI should exhibit an exponential decay across echo times. To examine deviations from this expected exponential decay, we calculated the voxel-wise percentage of unexplained variance. This calculation was performed by dividing the root mean square (RMS) of the residuals by the RMS of the observed signal, as shown in Figure S3. Regions with high susceptibility, such as the medial and lateral orbitofrontal cortices (OFC) and the entorhinal cortex, demonstrated a more pronounced nonexponential decay.

## 5 Supplementary Figures and Tables

**Figure S1.**
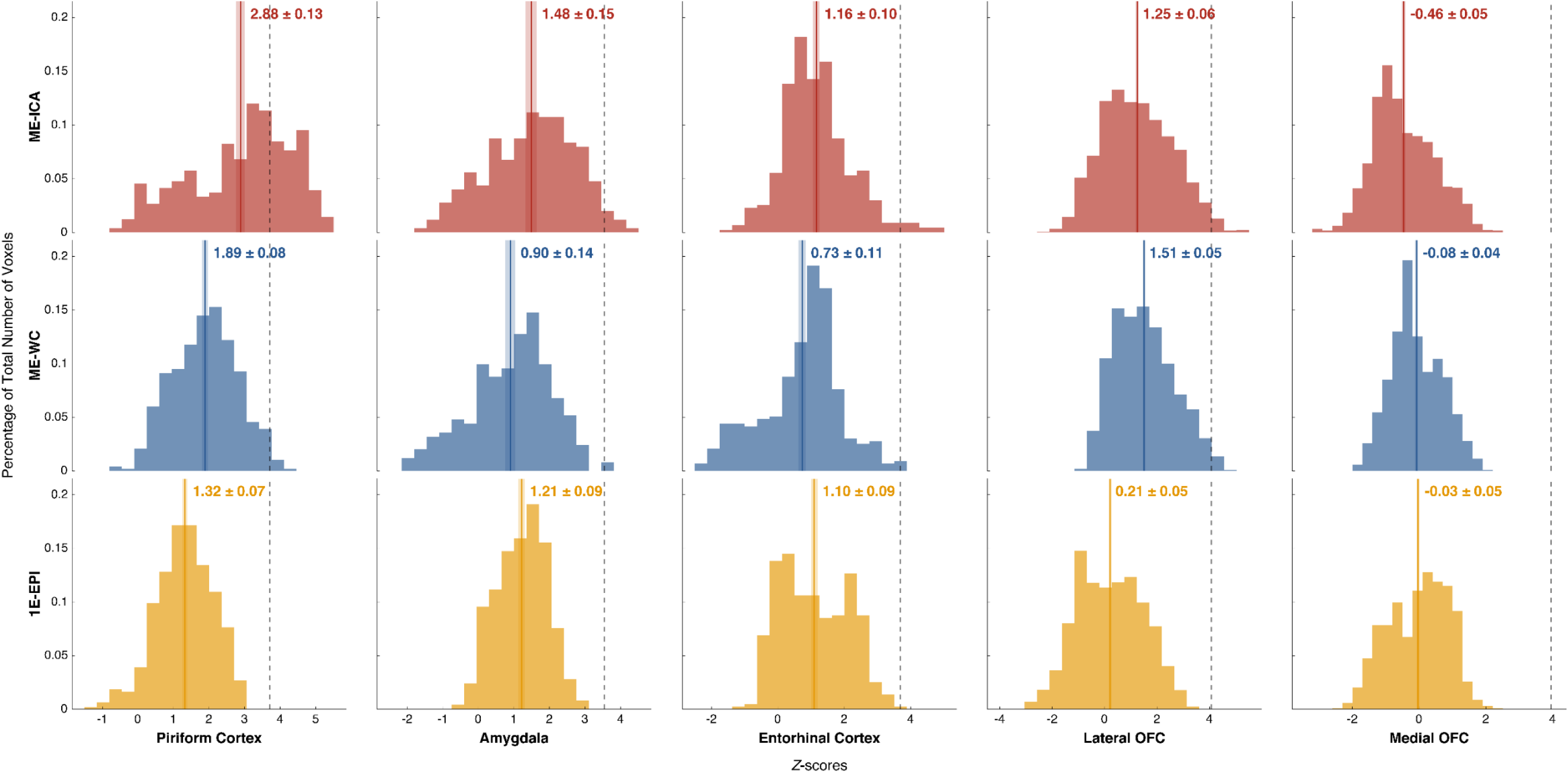
Histograms of Group Level Z-Scores for All ROIs for [Lemon > Control] Contrast. The mean ± 95% confidence interval of the Z-scores within each ROI for each acquisition is annotated. The mean is also marked by the solid line surrounded by a shaded area indicating the 95% confidence interval. The dashed line indicates the threshold corresponding to statistical significance (*p* < 0.05, SVC).

**Figure S2.**
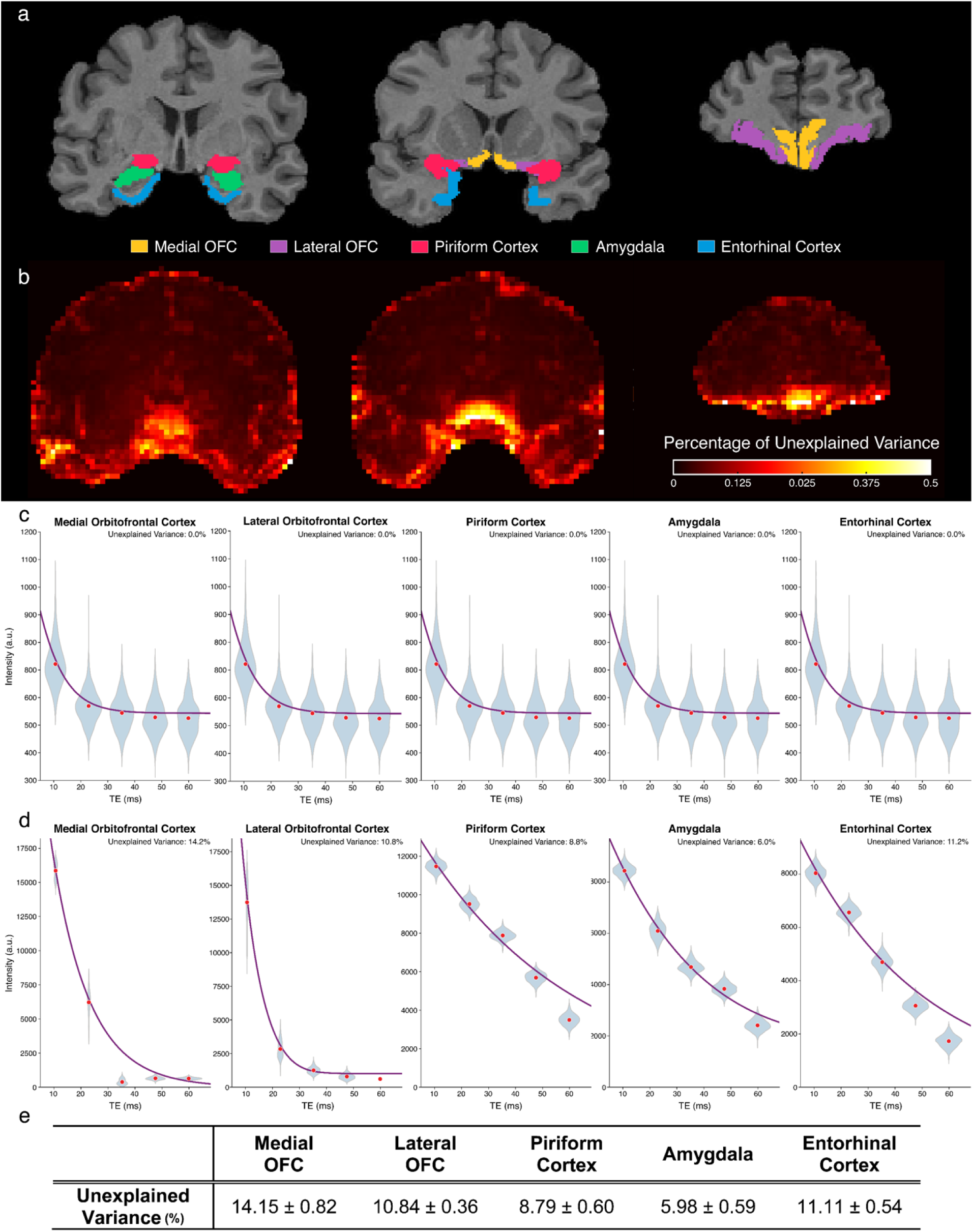
Percentage of Unexplained Variance in the Exponential Decay Signal across Different Echo Times for a Single Subject. a) Three representative coronal slices of a single subject with ROIs overlaid on T_1_-weighted anatomical images. b) The percentage of unexplained variance map for the representative coronal slices. c) Signal across different echo times for voxels with the minimum unexplained variance in each ROI. The red dots represent the mean signal intensity over 600 measurements (200 volumes per run, 3 runs in total), and the blue violin plots show the distribution of signal intensity. The purple lines indicate the fitted model using the method described in Section 2.7. d) Signal across different echo times for voxels with the unexplained variance closest to the mean unexplained variance within each ROI. The red dots, blue violin plots, and purple lines represent the mean signal intensity, distributions, and fitted model, respectively. e) The mean ± 95% confidence interval of unexplained variance for each ROI.

**Figure S3.**
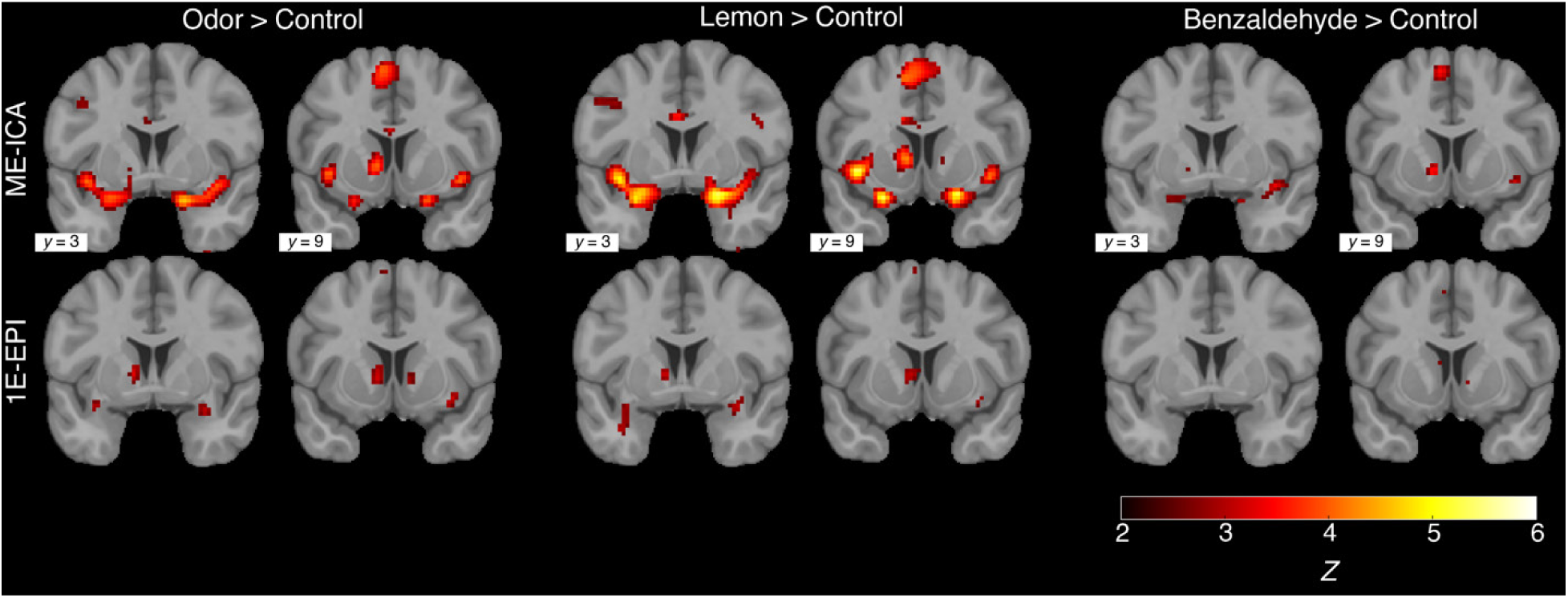
Comparison of Activation Maps among Different Contrasts for Olfactory-Related Tasks in Both 1E-EPI and ME-EPI. Two representative coronal slices of activation maps for all three contrasts ([Odor > Control], [Benzaldehyde > Control], [Lemon > Control]) using ME-EPI acquisition with ME-ICA denoising methods (ME-ICA) and using single-echo fMRI (1E-EPI) acquisition (*p* < 0.001, uncorrected).

**Figure S4.**
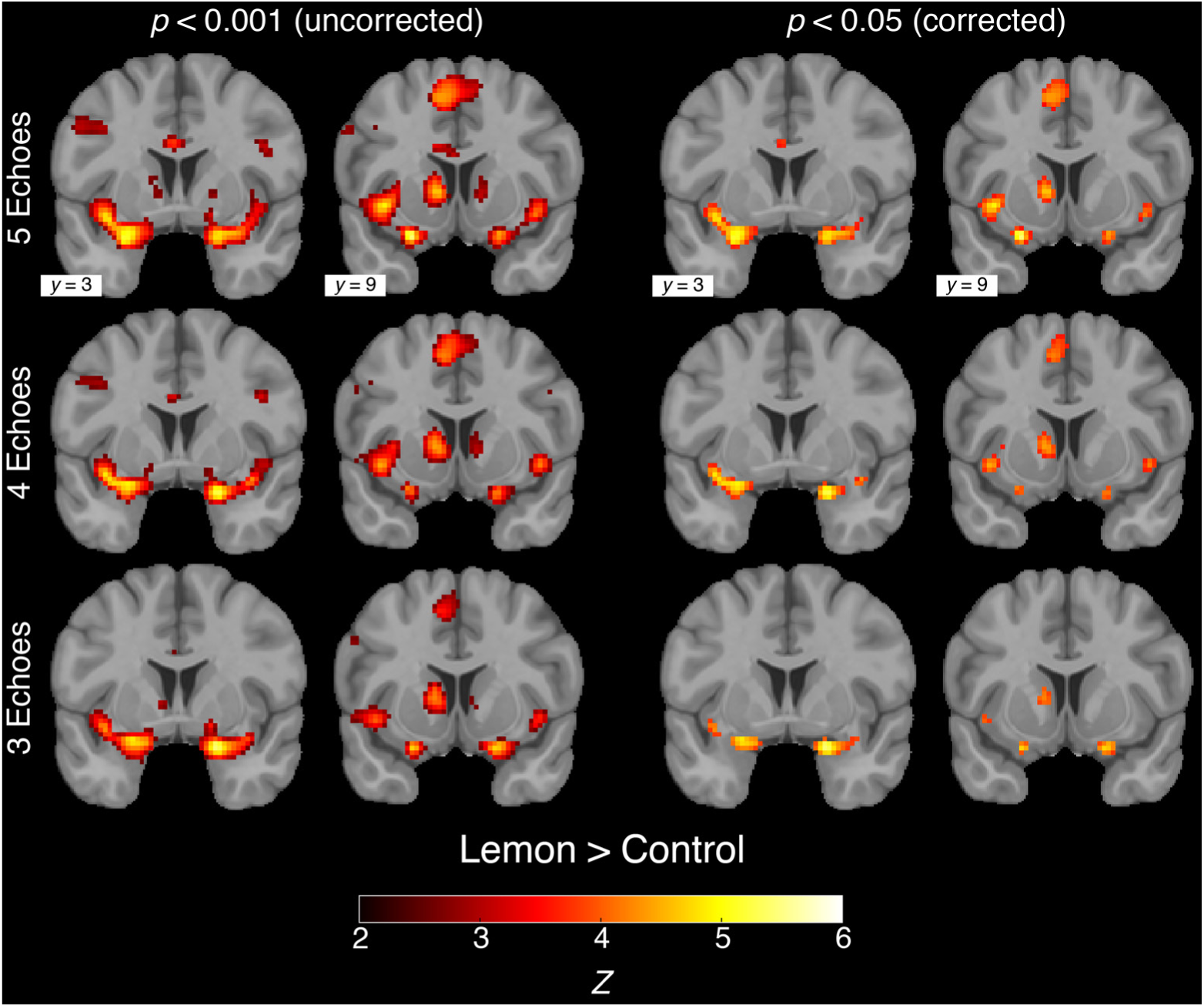
Comparison of Activation Maps for ME-EPI Acquisition with ME-ICA Denoising among Different Number of Echoes for Olfactory-Related Tasks. Two representative coronal slices of activation maps for [Lemon > Control] using ME-EPI acquisition with ME-ICA denoising methods (ME-ICA) with first 3, 4 and all 5 echoes. The left two columns show the maps without multiple comparison correction (*p* < 0.001) while the right two columns show the maps corrected for multiple comparison (*p* < 0.05, FDR corrected).

**Figure S5.**
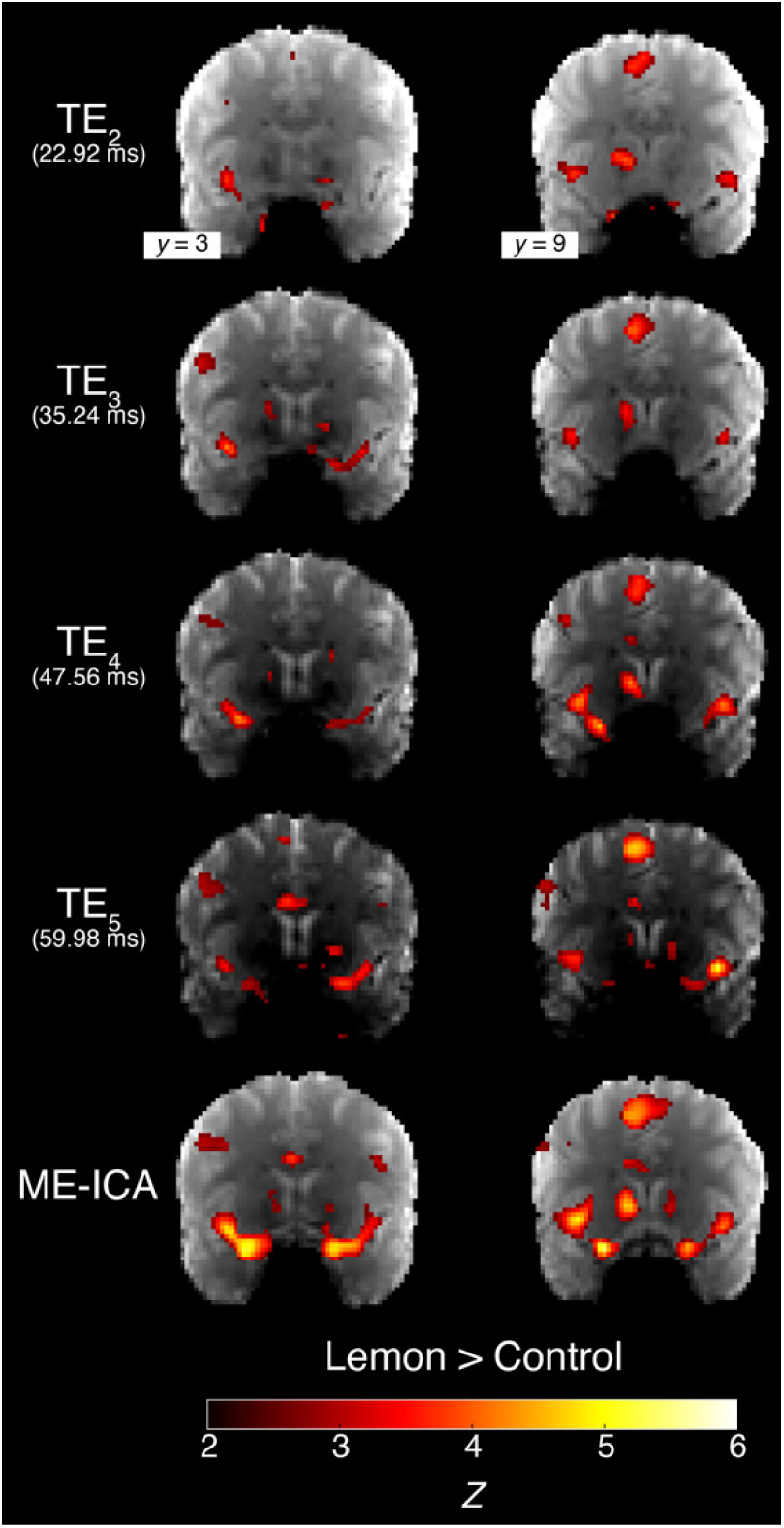
Comparison of Activation Maps When Analyzing Individual Echoes from ME-ICA Separately for Olfactory-Related Tasks. Two representative coronal slices of activation maps for [Lemon > Control] when analyzing individual echoes from ME-EPI acquisition with ME-ICA denoising methods (ME-ICA) separately on the group level (*p* < 0.05, uncorrected). The activation maps are overlaid on the mean image of a single subject across one run, registered to MNI template space, to illustrate signal dropout across different echoes.

**Table S1.**
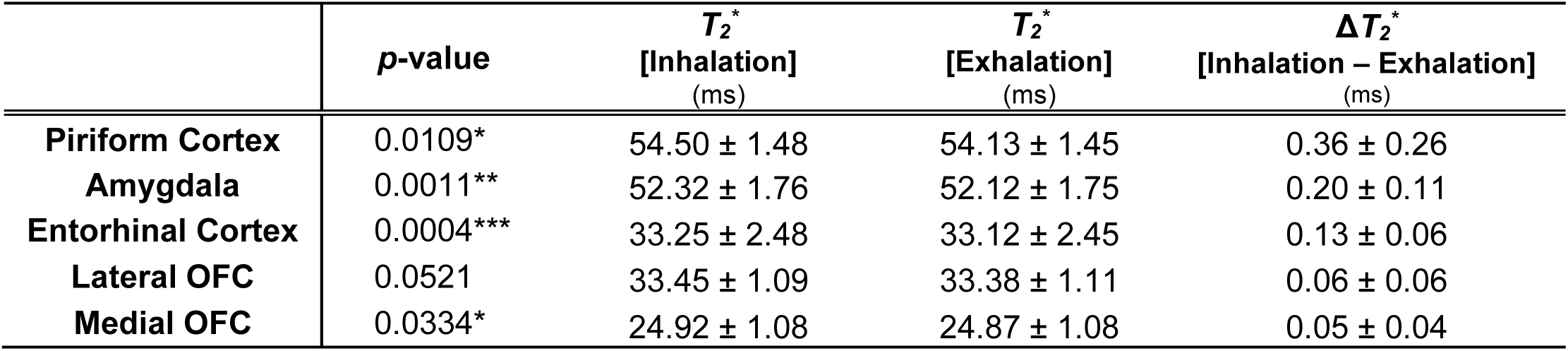
T_2_* Changes during Inhalation and Exhalation Phase in All Five ROIs. The *p*-values indicate the statistical significance of the two-sample, two-sided *t*-tests comparing T_2_ * values estimated from volumes during the inhalation phase with those from the exhalation phase. T_2_ * values and their differences (ΔT_2_*) are reported as mean values within the each ROI across all subjects ± 95% confidence intervals.

| | $p$-value | $T_2^*$<br>[Inhalation]<br>(ms) | $T_2^*$<br>[Exhalation]<br>(ms) | $\Delta T_2^*$<br>[Inhalation – Exhalation]<br>(ms) |
| --- | --- | --- | --- | --- |
| **Piriform Cortex** | 0.0109* | 54.50 ± 1.48 | 54.13 ± 1.45 | 0.36 ± 0.26 |
| **Amygdala** | 0.0011** | 52.32 ± 1.76 | 52.12 ± 1.75 | 0.20 ± 0.11 |
| **Entorhinal Cortex** | 0.0004*** | 33.25 ± 2.48 | 33.12 ± 2.45 | 0.13 ± 0.06 |
| **Lateral OFC** | 0.0521 | 33.45 ± 1.09 | 33.38 ± 1.11 | 0.06 ± 0.06 |
| **Medial OFC** | 0.0334* | 24.92 ± 1.08 | 24.87 ± 1.08 | 0.05 ± 0.04 |

**Table S2.** Z-Scores of ROI Analyses for Each Olfactory-related ROI under the Contrast [Lemon > Control] When Analyzing 5 Individual Echoes from ME-ICA Separately.

|  | Piriform Cortex | Amygdala | Entorhinal Cortex | Orbitofrontal Cortex |
| --- | --- | --- | --- | --- |
| **TE<sub>1</sub>**<br>(10.60 ms) | 1.945 | 1.010 | 0.756 | 1.205 |
| **TE<sub>2</sub>**<br>(22.92 ms) | 2.383 | 1.957 | 2.183 | 1.605 |
| **TE<sub>3</sub>**<br>(35.24 ms) | 3.457 | 1.583 | 1.789 | 1.724 |
| **TE<sub>4</sub>**<br>(47.56 ms) | 3.819 | 1.355 | 0.973 | 1.835 |
| **TE<sub>5</sub>**<br>(59.98 ms) | 3.693 | 2.604 | 1.265 | 1.705 |
| **1E-EPI**<br>(22.00 ms) | 2.479 | 1.972 | 2.152 | -0.279 |

**Table S3.** Comparison of Estimated T_2_* Values between Unmodified and Modified Fitting Methods for All ROIs within a Single Subject. Unmodified T_2_* estimation method fits Eq. (1) for each voxel. Modified T_2_* estimation method fits Eq. (3) instead.

|  | Medial OFC | Lateral OFC | Piriform Cortex | Amygdala | Entorhinal Cortex |
| --- | --- | --- | --- | --- | --- |
| **Unmodified $T_2^*$ Estimation**<br>(ms) | 21.9 ± 0.6 | 30.2 ± 0.6 | 45.8 ± 1.1 | 45.6 ± 1.1 | 29.0 ± 1.0 |
| **Modified $T_2^*$ Estimation**<br>(ms) | 13.0 ± 0.2 | 21.8 ± 0.2 | 41.3 ± 1.1 | 42.2 ± 1.0 | 22.2 ± 0.3 |

